# Spine loss in depression impairs dendritic signal integration in human cortical microcircuit models

**DOI:** 10.1101/2024.06.19.599729

**Authors:** Heng Kang Yao, Frank Mazza, Thomas Prevot, Etienne Sibille, Etay Hay

## Abstract

Major depressive disorder (depression) is associated with altered dendritic structure and function in excitatory cortical pyramidal neurons, due to decreased inhibition from somatostatin interneurons and loss of spines and associated synapses, as indicated in postmortem human studies. Dendrites play an important role in signal processing as they receive the majority of synaptic inputs and exhibit nonlinear properties including backpropagating action potentials and dendritic Na^+^ spikes that enhance the computational power of the neuron. However, it is currently unclear how depression-related dendritic changes impact the integration of signals. Here, we expanded our previous data-driven detailed computational models of human cortical microcircuits in health and depression to include active dendritic properties that enable backpropagating action potentials as measured in human neurons, and spine loss in depression in terms of synapse loss and altered intrinsic property. We show that spine loss dampens signal response and thus results in a larger impairment of cortical function such as signal detection than due to reduced somatostatin interneuron inhibition alone. We further show that the altered intrinsic properties due to spine loss abolish nonlinear dendritic integration of signals and impair recurrent microcircuit activity. Our study thus mechanistically links cellular changes in depression to impaired dendritic processing in human cortical microcircuits.

## Introduction

Major depressive disorder (depression) is associated with an altered cortical excitation-inhibition balance that disrupts signal processing^1,2^. Recent studies in humans and in chronically-stressed rodents implicated altered dendritic mechanisms in cortical pyramidal (Pyr) neurons, due to reduced apical dendritic inhibition from somatostatin (SST) interneurons^1,3^ and loss of dendritic spines and synapses^4–7^. We previously used detailed models of the human cortical microcircuits to link reduced SST interneuron inhibition in depression to functional impairments in signal detection^8^. However, the effects on dendritic input processing and the involvement of spine loss remain unclear.

Reduced SST interneuron inhibition in depression is supported by reduced expression of SST and other GABAergic markers of SST interneurons in postmortem brain tissue from depression patients^3,9^. Similar changes in expression are also seen in chronic stress in rodents, which exhibit depressive and cognitive symptoms^10^. In support of a causal link, brain-wide silencing SST interneurons in rodents produced depression symptoms and cognitive impairments^11^. Chronic stress in rodents also induces a loss of spines in the basal and apical dendrites of Pyr neurons^6^, and a loss of synapses on spines in the dorsolateral prefrontal cortex was also found postmortem in depression patients^7^. Accordingly, application of pharmacology modulating α5-GABA_A_ receptors that SST interneurons target had antidepressant and pro-cognitive effects and also increased spine density in chronic stress in rodents^6,12^.

Dendrites contribute considerably to the computational power of neurons and microcircuits, by receiving and transforming signals through nonlinear properties^13–16^. Dendritic spines in excitatory Pyr neurons receive the majority of excitatory synaptic input^17^ and also a considerable portion of inhibitory synaptic input^18^. The spines increase the surface area of the dendrites and thus the number of signals that can be integrated, and also affect the passive conductance and membrane capacitance and thus the excitability of the neuron^19–21^. Dendritic Na^+^ channels allow for effective back-propagated action potentials (bAP) from the soma to the distal dendrites^22^, which summate with incoming synapses to enhance dendritic processing and coincidence detection^13,23^. Dendritic ion channels also mediate local dendritic Na^+^ spikes^22^ that enable local dendritic non-linear computations, enhancing the neuron’s computational power^24^. bAPs and dendritic Na^+^ spikes can be inhibited by GABAergic inputs and therefore may be impacted by reduced inhibition and loss of synapses in depression^25,26^.

While rodent studies have linked reduced inhibition and spine loss to cognitive impairments^6^, a mechanistic link remains to be established, particularly in the human context. Nonlinear processing may differ between rodents and humans, as human dendrites are more complex and compartmentalized, and have larger diameters that result in higher thresholds for NMDA spikes^14,27^. In addition, human synapses are stronger with lower failure rates^28–31^. As such, the loss of synapses due to spine loss in depression may have larger or different effects in humans. We previously integrated human neuronal, synaptic, and connectivity data to generate detailed biophysical models of human cortical microcircuits in health and depression, and used them to link reduced SST interneuron inhibition with effects on cortical microcircuit activity and function^8^. Recent studies characterized dendritic bAP in human Pyr neurons^13^, providing important constraints to better simulate dendritic processing in health and depression.

In this study, we expanded our previous models to include human dendritic bAP properties, and spine loss in depression. Using these new models, we studied the individual and combined effects of reduced SST interneuron inhibition and spine loss on cortical microcircuit baseline activity, signal response and signal detection error rates. Furthermore, we studied how changes in intrinsic properties due to spine loss can impact bAP-mediated dendritic processing on the cellular and microcircuit level.

## Results

To investigate the impact of depression mechanisms on dendritic processing in human microcircuits, we first expanded our previous model of human cortical L2/3 Pyr neuron to include active dendrites with bAP. We uniformly distributed K^+^ and Na^+^ ion channels (Na_T_: 0.0097 S/cm^2^, K_V3_1_: 0.101 S/cm^2^) on the apical dendrites, and generated models with bAP amplitudes similar to those recorded in human dendrites (Fig. 1A). Compared to the passive dendrite, the active dendrites amplified the bAP amplitude by 194.5% ± 42.8%. In addition, the refitting of the Pyr models with active dendrites maintained a similar performance across all other electrophysiological features (step response, sag) as our previous model. We then integrated the new Pyr neuron models into our previous models of human L2/3 cortical microcircuits, and calibrated connection probabilities and background Ornstein-Uhlenbeck (OU) conductance^32^ to maintain the fit of average baseline firing rate and response profile in the different neuron types (Fig. 1B-D). The new models with active dendrites also maintained the prominent alpha peak in the simulated electroencephalogram (EEG) power spectral density (PSD) and improved on the previous models by having higher power in low-theta frequencies (4 - 6 Hz), thus better capturing the 1/f relationship seen in human resting-state EEG (Fig. 1E).

**Fig 1.**
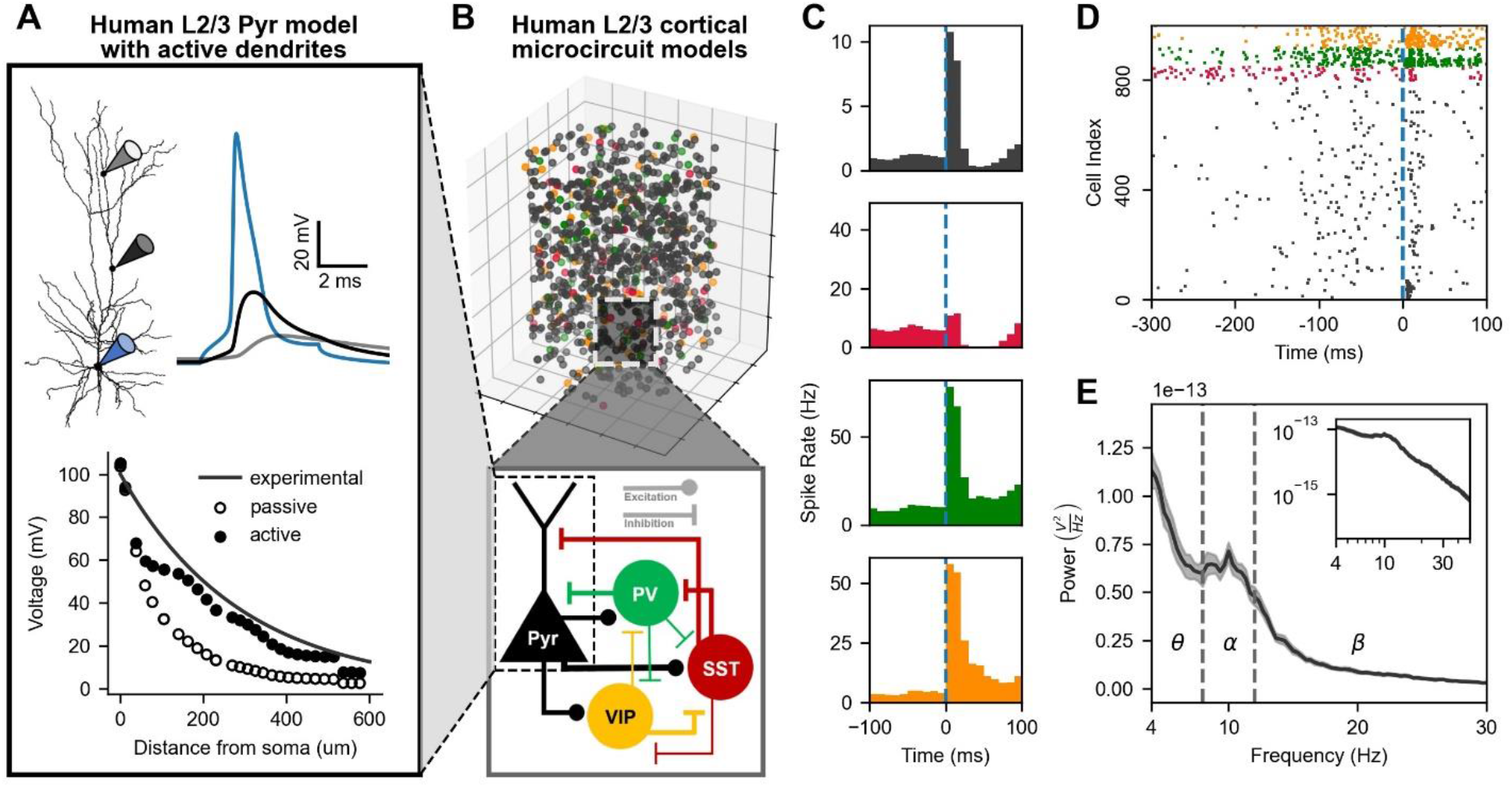
Human L2/3 cortical microcircuit models with active dendrites. **A**. Model of human cortical L2/3 Pyr neuron with active dendrites and bAP. Top: human Pyr neuron morphology and simulated voltage at the soma (blue), 200 μm from soma (black) and 400 μm from soma (gray). Bottom: amplitude of bAP with distance from the soma in models with active and passive (Na^+^ and K^+^ channels blocked) dendrites. Solid black line shows the experimental fit from the literature. **B**. Models of human L2/3 cortical microcircuits included 1000 neurons of the four key types (800 Pyr, 50 SST, 70 PV, 80 VIP) distributed in a 500×500×950 μm^3^ volume according to human dimensions. Bottom – schematic of the microcircuit connectivity between the different neuron types. **C**. Average peristimulus time histogram of microcircuit activity (n = 100 simulated microcircuits). Vertical dashed line signifies onset of a brief stimulus (∼5 ms). **D**. Example raster plot of microcircuit activity at baseline and response. Vertical dashed line signifies the stimulus onset. **E**. PSD of simulated EEG from the microcircuit (n = 10 microcircuits). Dash lines separate different frequency bands. Inset shows the plot in log-log scale.

We used the models of human L2/3 cortical microcircuits with active dendrites to generate depression microcircuit models that included two key altered mechanisms: 40% reduced SST interneuron inhibition (SST↓), 30% spine loss in Pyr neurons (spine↓), and their combined effects (SST↓/spine↓, Fig. 2A). Critically, spine loss involved the loss of excitatory and inhibitory synapses, as well as membrane surface area. Consistent with our previous findings, reduced SST inhibition in the expanded models significantly increased baseline activity (healthy: 0.72 ± 0.03 Hz; SST↓: 1.21 ± 0.03 Hz, 68% increase, *p* < 0.05, Cohen’s *d* = 13.9, Fig. 2B) but had a negligible effect on the response (healthy: 2.9 ± 0.44 Hz; SST↓: 2.82 ± 0.39 Hz, 3% decrease, *p* < 0.05, Cohen’s *d* = 0.39, Fig. 2B). Spine loss increased baseline activity as well, but to a lower extent (1.03 ± 0.02 Hz, 43% increase, *p* < 0.05, Cohen’s *d* = 12.2). However, unlike SST interneuron inhibition loss, spine loss decreased the response rates (2.03 ± 0.15 Hz, 30% decrease, *p* < 0.05, Cohen’s *d* = 2.9, Fig. 2B-D), primarily by decreasing recurrent activity that followed the bottom-up activation (∼10 - 30 ms post-stimulus, Fig. 2D). When both mechanisms were combined, the effect on baseline activity was slightly higher than SST↓, indicating a sublinear summed effect (1.33 ± 0.03 Hz, 85% increase, *p* < 0.05, Cohen’s *d* = 23.9), whereas the response rates were decreased to an extent similar to the condition with only spine loss (1.99 ± 0.15 Hz, 31% increase, *p* < 0.05, Cohen’s *d* = 2.9, Fig. 2B-D).

**Fig 2.**
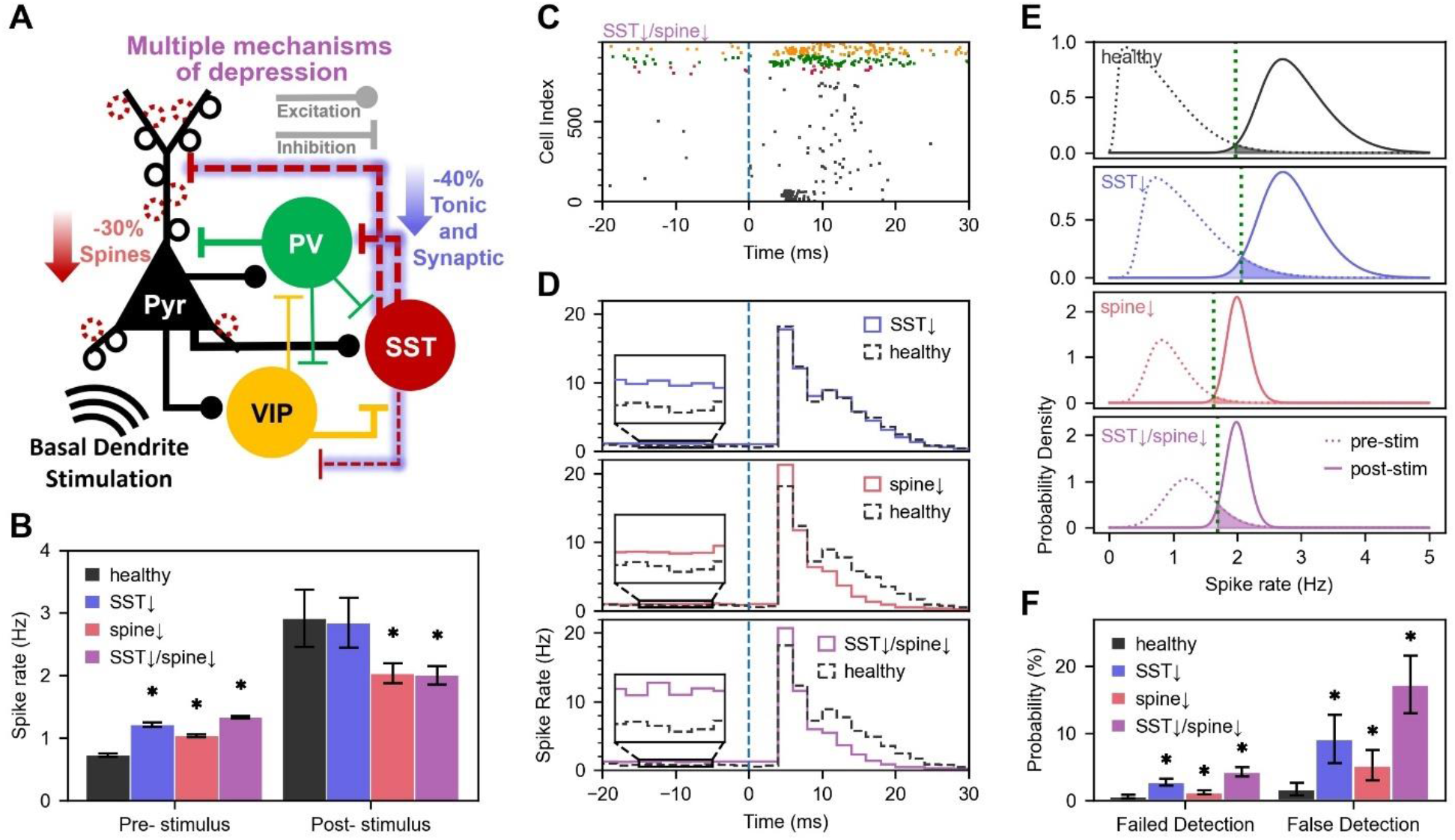
Spine loss in depression dampens response to stimuli. **A**. Schematic of depression models for reduced SST inhibition (SST↓) and spine loss (spine↓). **B**. Average baseline and response spike rate of Pyr neurons in healthy and depression microcircuits. **C**. Example raster plot of SST↓/spine↓ microcircuit activity at baseline and response. Vertical dashed line signifies the onset of the stimulus. **D**. Average peristimulus time histogram of microcircuit activity in healthy vs. depression conditions (n = 100 simulated microcircuits per condition) **E**. Distribution of pre-stimulus firing rates in 50-ms windows in example healthy and depression microcircuits (n = 1,950 windows each) and distribution of firing rates 50 ms post-stimulus across 100 microcircuits (n = 100 windows, bootstrapped 1000 times). The vertical dotted line denotes the decision threshold of signal detection. **F**. Summary of false and failed stimulus detection rates in healthy and depression microcircuits (n = 100 microcircuits per condition). Asterisk denotes p < 0.05 and large effect size. Data are represented as mean ± SD.

To investigate the effects of depression mechanisms on signal detection error rates, we calculated the failed and false detection rates from the distribution of Pyr firing rate in 50 ms windows at baseline and during response (Fig. 2E). In the SST↓ condition, the distribution of baseline rate was right-shifted, resulting in increased failed detection (healthy: 0.58 ± 0.2%, SST↓: 2.69 ± 0.53%, *p* < 0.05, Cohen’s *d* = 5.0, Fig. 2F) and false detection (healthy: 1.70 ± 0.94%, SST↓: 9.15 ± 3.59%, *p* < 0.05, Cohen’s *d* = 2.8). In the spine↓ condition, there was a considerable left shift of the response rate distribution, and also a narrowing of both baseline and response distributions, which resulted in an overall increase in failed detection (1.18 ± 0.31%, *p* < 0.05, Cohen’s *d* = 2.1) and false detection (5.21 ± 2.25%, *p* < 0.05, Cohen’s *d* = 2.0), but to a lesser extent than in the SST↓ condition. In the case of SST↓ and spine↓ combined, the effects summated supra-linearly for both failed detection (spine↓/SST↓: 4.25 ± 0.71%, *p* < 0.05, Cohen’s *d* = 6.9, compared to healthy; SST↓ only + spine↓ only: 3.87 ± 0.61%, *p* < 0.05, Cohen’s *d* = 0.57, compared to spine↓/SST↓) and false detection (spine↓/SST↓: 17.29 ± 4.31%, *p* < 0.05, Cohen’s *d* = 5.0, compared to healthy; SST↓ only + spine↓ only: 14.36 ± 4.24%, *p* < 0.05, Cohen’s *d* = 0.68, compared to spine↓/SST↓) due to a combined right shift of the baseline rate distribution and left shift of the response spike rate distribution.

After characterizing the effect of the depression mechanisms on the microcircuit response to bottom-up inputs, we turned to investigate the effect of spine loss on dendritic integration with bAP (Fig. 3A). The decreased membrane capacitance and leak conductance due to spine loss in depression resulted in abnormally large bAP amplitude (44.3 ± 10.9 mV; 261.6 ± 108.5% Fig. 3B) in 3 of the 6 major paths from midway of the dendritic tree to the distal end (> 300 μm from the soma). These abnormally large bAPs also had decreased half-width (-5.4 ± 1.6 ms; -72.3 ± 12.1% Fig. 3C). To investigate how these changes in bAP amplitude and half-width impact the integration of incoming dendritic synaptic inputs, we elicited PSPs of increasing strength at 400 μm from the soma shortly after the peak of the bAP (6 ms after the somatic stimulation) and calculated the integral of the local dendritic voltage between 6 and 12 ms post-stimulus (Fig. 3D). In the healthy condition, PSPs integrated nonlinearly with the bAP, exhibiting a sharp increase in dendritic depolarization beyond a certain magnitude of synaptic input. The non-linearity was abolished in the spine↓ condition (dendritic voltage integral for 0.0024 μS synaptic conductance relative to no synaptic input: healthy: 137.0 mV*ms, spine↓: 91 mV*ms, 33.6% decrease, Fig. 3E). To investigate whether the nonlinear dendritic integration impacted the soma, we measured the peak voltage in the soma following the dendritic stimulation (between 10 and 14 ms post-stimulus). We similarly found a nonlinear increase in depolarization that was present in the healthy condition but not in the spine↓ condition, resulting in greater somatic excitation in the healthy neuron (somatic voltage for 0.0024 μS synaptic conductance relative to no synaptic input: healthy: 0.83 mV, spine↓: 0.54 mV, 35% decrease, Fig. 3F).

**Fig 3.**
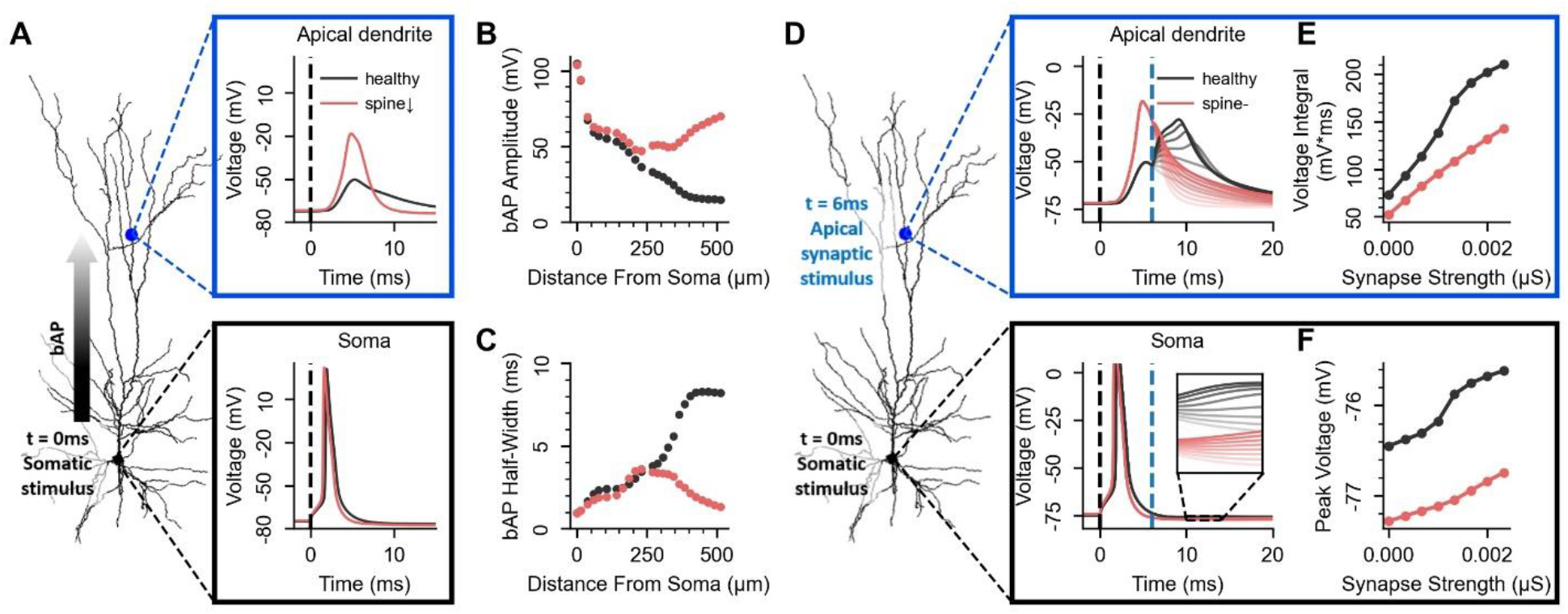
Spine loss in depression impairs nonlinear dendritic integration with bAP. **A**. Simulation of a bAP travelling along the apical dendrite in a Pyr neuron in health (black) and depression (spine↓, red), showing voltage traces at distal apical dendrites (400 μm from soma, top) and at the soma (bottom). **B**. Amplitude of bAP at different distances from the soma. **C**. Half-width of bAP at different distances from the soma. **D**. Simulation of a bAP travelling along the apical dendrite in a Pyr neuron, coincident with incoming apical synaptic inputs of increasing strength, in health and depression. Plots show the voltage traces at the location of synaptic stimulation (400 μm from soma, top) and at the soma (bottom), with the opacity of the colours corresponding to the synapse strength. The inset shows the somatic voltage traces at 10 - 14 ms post-stimulus, analyzed in panel F. **E**. Integral of dendritic voltage over 6 - 12 ms post-stimulus in response to different synaptic strength, in health and depression. **F**. Peak voltage in the 10 - 14 ms window post-stimulus at the soma, in health and depression.

We next investigated the effect of spine loss on dendritic input integration in the microcircuit by stimulating a population of 55 Pyr neurons with a similar protocol as above, with a suprathreshold somatic stimulus (t = 0 ms) and synaptic input at the distal apical dendrite (t = 6 ms) with varying activation strength (0 - 100 synapses, Fig. 4A). The apical inputs generated a second peak in the peristimulus time histogram (PSTH) spiking response at around 10 ms after the somatic stimulation, and increased recurrent activity from the time of dendritic stimulation (6 ms) to 30 ms post-stimulus (Fig. 4B, C). Across all numbers of dendritic synapses that we have simulated, depression microcircuits had reduced firing rates compared to healthy microcircuits (for example, 100 synapses: SST↓/spine↓: 5.81 Hz ± 0.91 Hz, healthy: 10.7 Hz ± 1.95 Hz, *p* < 0.05, Cohen’s *d* = 3.2, Fig. 4D). Moreover, the gain of the response rate as a function of the number of apical synaptic inputs was reduced in depression (SST↓/spine↓: 4.15 ± 0.38 Hz/100 synapses, healthy: 8.77 ± 1.37 Hz/100 synapses, *p* < 0.05, Cohen’s *d* = 4.6, Fig. 4D).

**Fig 4.**
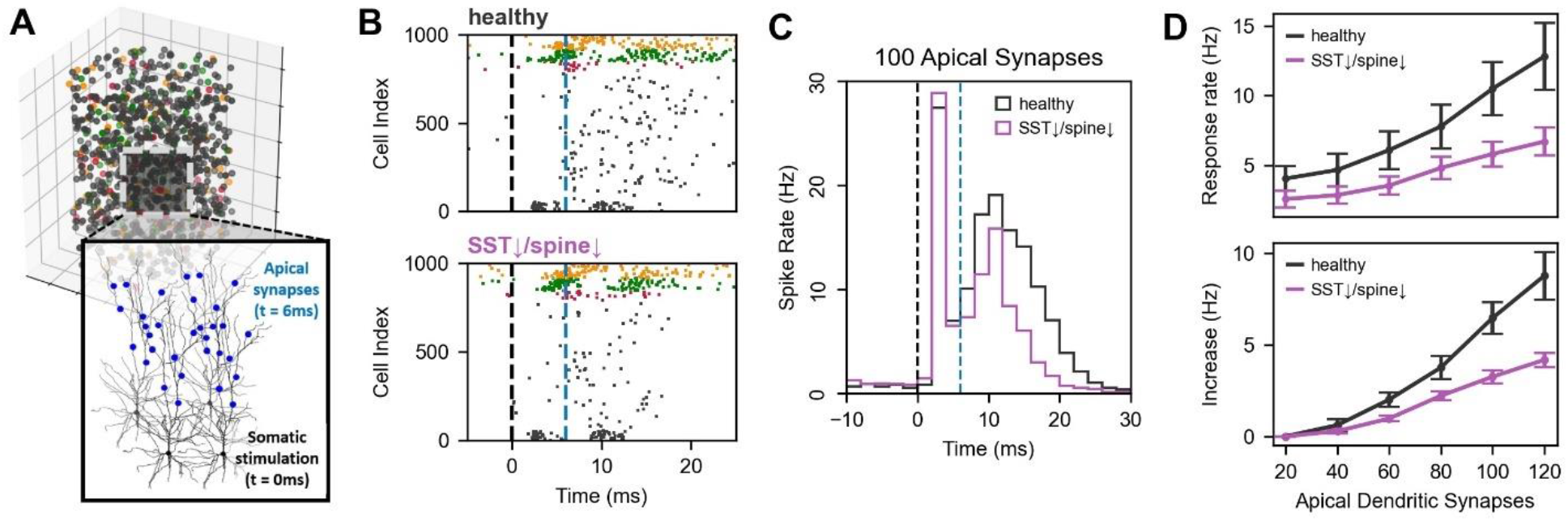
Decreased nonlinear integration of signals in depression microcircuits. **A**. Schematic showing stimulation protocol of a microcircuit, where 55 Pyr neurons were stimulated at the soma, followed (t = 6 ms) by stimulation at the distal apical dendrites with varying activation strength (0 - 120 synapses). **B**. Example raster plots showing baseline and response to 100 apical dendritic synaptic inputs in healthy and depression. Black and blue vertical dashed line show the start of somatic and dendritic stimulation, respectively. **C**. PSTH of baseline and response to somatic and distal apical stimulation with 100 synapses. **D**. Top - bootstrapped mean spike rate of Pyr neurons in response to somatic and apical synaptic stimulation, in healthy and depression microcircuits. Bottom - increase in mean spike rates relative to the case of 20 apical dendritic synapses, in healthy and depression microcircuits.

## Discussion

In this study, we linked altered dendritic mechanisms in depression with impaired signal integration in human neurons and cortical microcircuits. To overcome limitations of studying dendrites and microcircuits in the living human brain, we expanded our previous detailed computational models to reproduce human active dendritic properties of bAP and dendritic spine loss in depression. We demonstrated that spine loss impaired response firing, in contrast to reduced SST interneuron inhibition mainly affecting baseline firing. Furthermore, we showed that spine loss abolishes the non-linear synaptic integration mediated by bAP and thus impairs cortical recurrent activity. Our results with the expanded models differentiate effects of reduced SST interneuron inhibition and spine loss on dendritic signal processing in depression. These new models can be used to refine corresponding EEG biomarkers^33^ and enable more accurate estimation of the effects of novel depression pharmacology^34^ that targets the apical dendrites.

Our findings of reduced response due to spine loss in depression can explain reduced cortical activity in patients during somatosensory response^35^ and in cognitive tasks^36–38^, which was correlated with depression severity^35^. The decreased recurrent activity due to spine loss that we found could play a role in the reduced sustained neural activity in depression patients when processing positive emotional stimuli^39^. Our results may extend to diseases such as Alzheimer’s disease^40^ and schizophrenia^41^, where spine loss and reduced cortical activity during response^42^ are also observed and are correlated with cognitive deficits^43^. We modeled spine loss according to rodent data in mild chronic stress, which is of a similar order of magnitude to the level of spine synapse loss in post-mortem human tissues^6,7^ although somewhat lower. A possible reason is that mild chronic stress is more similar to an early or induction stage of depression, with moderate spine loss^44^. Another reason may be that the human study^7^ involved older subjects, with potentially fewer spines due to aging^45^. We used the rodent data as it was richer and enabled better constraints for spine loss in basal and apical dendrites and did not have the age confound that the human study had. We modelled spine loss as decreased number of synaptic contact points, without affecting connection probability. While the connectivity changes underlying synapse loss are unclear and can alternatively be modelled as decreased connection probability between neuron types, our choice corresponded more directly to the available data. The net loss of synapses and change in the excitation/inhibition should be similar in either case, but future work can compare the effects of the different implementations. We modeled spine loss in terms of altered intrinsic properties of dendrites and did not model spines in detail, since the change in intrinsic properties sufficed to investigate the effects on overall dendritic integration, but future studies can use more detailed models of dendritic spines^46^ to study how reduced spine density in depression impacts localized computations within spines^47^. Future studies can also examine the effects of reduced dendrite length and complexity, which were observed in restraint stress rodent model studies and suggested by reduced microtubule-associated protein 2 in post-mortem human tissue in depression patients^7^, a marker associated with dendrite structure. However, these changes remain out of the scope of the current study due to lack of sufficient human data.

The abolishment of non-linear dendritic integration in spine loss resulted from abnormally large amplitudes of bAP, which shortened the dendritic bAP half-width and thus the summation time window with PSPs, and also deactivated sodium channels that mediate nonlinear boosting of PSP^48,49^. Similar large amplitude bAP has been observed in rodents due to changes in dendritic sodium and potassium channels^50^, and here we demonstrate an alternative mechanism of decreased membrane capacitance and passive conductance due to spine loss. bAP has also been shown to mediate dendritic release of BDNF^51^, a key transcription factor that is reduced in depression and plays a role in maintaining dendritic spines and SST expression^52–54^. Altered bAP in depression may therefore have further effects beyond the ones we have simulated. Our models reproduced bAP amplitudes, but the propagation of bAP was slower in our models than measured experimental values^13^, possibly due to the dendrites being more branched in the morphology we used^55^. However, the difference would mainly shift the relevant temporal delay of bAP and PSP summation, for which our spine loss results would hold as well. Future studies can look into using different morphologies to improve the propagation speed. We did not model bAP in the basal dendrites due to a lack of corresponding human data. We also did not model dendritic calcium spikes, which were shown to uniquely enable anti-coincidence detection of signals in humans^13^, because data of the ionic channel mechanisms by which these calcium spikes arise is not yet available. Our expanded models with active dendrites can improve identifying corresponding *in-silico* EEG biomarkers^33^, since bAP and dendritic currents significantly contribute to EEG signal^56,57^. Our models can also be used to improve the *in-silico* testing of novel pharmacology that targets SST interneuron inhibition pathway^34^, by estimating the pharmacology effects on dendritic integration.

## Methods

### Human L2/3 Pyr neuron models with active dendrites

Following similar methods as in our previous work^8^, we generated multicompartmental conductance-based single neuron models for Pyr neurons with active dendrites using BluePyOpt^58^. We refitted the firing and passive electrophysiological features as our previous work^8^ together with bAP amplitude features. To fit the bAP in the Pyr neuron, we injected a 1.0 mA current at the soma for 5 ms to elicit somatic action potentials and fitted the bAP amplitude along the apical dendrite at distances of 200 μm and 400 μm from the soma according to literature values (200 μm: ∼60mV, 400 μm: ∼15mV)^13^. We distributed Na_T_ and K_v3_1_ channels uniformly in the apical dendrites. I_h_ channel was distributed with a sigmoidal function with the absolute distance from the soma:

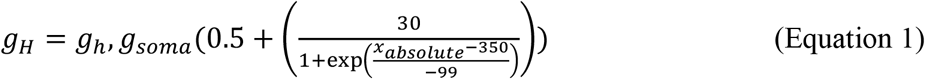

Apical dendritic Na_T_ kinetics parameters were the same as the axonal parameters based on past work^59,60^ (Vshift_m_ = 0, Vshift_h_ = 10, Slope_m_ = 9, and Slope_h_ = 6). We used the same Pyr neuron morphology as our previous models, from the Allen Brain Atlas^61^. To run the model optimization, we used parallel computing clusters on SciNet^62^ with 280 processors. The optimization used a population size of 280 over 400 generations for an approximate total runtime of 6 hours.

### Human L2/3 microcircuit models

We integrated the new Pyr neuron model into our previous models of human cortical L2/3 microcircuits^8^, comprised of 1000 neurons (800 Pyr, 50 SST, 70 PV, and 80 VIP) distributed within a 500×500×950 μm^3^ volume (250 to 1200 μm below pia, spanning L2/3^63^). The microcircuit models included AMPA/NMDA and GABA_A_ synaptic mechanisms^15,64,65^, tonic inhibition^66^, and were driven by random OU background inputs^32^. The models were constrained with human data of neuronal firing and passive properties for each of the neuron types, synaptic properties for most of the connection types, cell proportions, and *in vivo* baseline firing rate in Pyr neurons, where synaptic and connectivity properties for which human data was not available were constrained with primate or rodent data (see details in Yao et al. 2022, Table S1).

PV and VIP interneuron models remained unchanged from our previous models, while SST interneuron models were refitted to follow the same methodology as the rest of the neuron models, with single optimization of all features and using the BluePyOpt tool^58^. Synaptic connections targeting the new Pyr and SST neuron models were refitted to reproduce similar average amplitude as our previous models^8^. Tonic inhibition was re-estimated with the new Pyr neuron model with active dendrites following the same methods as previously^8,67,68^, resulting in a similar value (previous: 0.938 mS/cm^2^, current: 0.994 mS/cm^2^). We refitted the resulting microcircuits with new Pyr neuron and SST interneuron models to reproduce average *in vivo* firing rates across cell types^69–71^ during baseline as in our previous models^8^, by adjusting the microcircuit connection probability and synaptic conductance, and the strengths of OU background inputs (Table 1). To reproduce average spike rates and dynamics during response to a brief stimulus as in our previous models, we stimulated 70 Pyr neurons with 7 basal dendritic synapses (t: 2 - 3ms, g: 3 μS), 30 PV interneurons were with 10 dendritic synapses (t: 0 – 3 ms, g: 3 μS) and 45 VIP interneurons were with 10 dendritic synapses (t: 0 – 6 ms, g: 3 μS). Microcircuit models were simulated using NEURON^72^ and LFPy^73^.

**Table 1.** Parameter changes relative to the previous model. Orange and green signify a decrease and increase, respectively, compared to the previous microcircuit model^8^. For synaptic conductance and connection probability, rows correspond to the presynaptic neuron and columns correspond to the postsynaptic neuron.

| Synaptic conductance (pS) | PYR | SST | PV | VIP |
| --- | --- | --- | --- | --- |
| PYR | 0.0000848 | -0.000044 | 0 | 0.00002 |
| SST | 0.000416 | 0.00008 | 0.00012 | 0 |
| PV | 0.00070166 | 0.00009 | 0.00012 | 0.00008 |
| VIP | 0 | 0.00006 | 0.00011 | 0.00008 |
| Connection probability (%) | PYR | SST | PV | VIP |
| PYR | 0 | -0.05 | 4.3 | 2 |
| SST | 0 | 0 | 6 | 12 |
| PV | 1.6 | -1 | -3 | 0 |
| VIP | 0 | -10 | 4 | -2 |
| OU conductance (pS) | 0.0000023 | 0.0000117 | -0.000078 | -0.0000012 |

### Simulated EEG

Using LFPy, we computed dipole moment from transmembrane currents within each cell in the microcircuit and generated the EEG timeseries using a four sphere volume conductor model with an EEG electrode placed directly above the circuit^33,73^. EEG PSD was calculated using Welch’s method with a Hanning window of 3 seconds^74^.

### Simulating coincidence of bAP and synaptic input

We stimulated the soma with a step current of 0.7 mA for 2 ms to elicit a somatic action potential, and 6 ms after the somatic stimulation we followed by stimulation of a synapse on the apical dendrite (400 μm from the soma) with varying conductance strength (0 – 0.0024 μS). As a corresponding protocol of coincident somatic and apical inputs in the microcircuit simulations, we stimulated 55 Pyr neurons with 7 somatic synapses (t: 0 – 1 ms, g: 3 μS) and 20 - 120 distal apical dendritic synapses (t: 6 – 7 ms, g: 0.3 μS). To balance the microcircuit response, as in the simulations of brief stimulus response, we also stimulated 20 PV neurons with 10 dendritic synapses (t: 0 – 3 ms, g: 3 μS), and 38 VIP neurons with 10 dendritic synapses (t: 0 – 5 ms, g: 3 μS).

### Reduced SST interneuron inhibition depression microcircuit models

We used our previous models of depression microcircuits^8^, with 40% decreased SST interneuron synaptic and tonic inhibition, based on expression studies in post-mortem brain tissue of depression patients^3^. When decreasing the tonic inhibition, the calculation of the relative contribution of each interneuron type to the total tonic inhibitory input was now refined to also account for the average firing rates of each interneuron type.

### Spine loss depression microcircuit models

We modelled 30% spine density loss in the apical and basal dendrites of Pyr neurons, according to studies in rodents under chronic stress and in post-mortem human tissue^6,7^, by reducing the membrane capacitance and passive conductance due to spines in the apical and basal dendrites by 30% (equivalent to 15% of the total dendritic membrane capacitance and passive conductance, assuming a doubling of the surface area. i.e. an additional 1 μf/cm^2^, for spine membrane area compensation^8^). To account for the loss of synapses due to spine loss, we reduced the number of synaptic contact points in connections that target Pyr neurons: as 80% of excitatory synapses and 30% of inhibitory synapses target spines^17,18^, we reduced contact points in Pyr→Pyr connections by ∼24% (from 3 to 2), and inhibitory synapses onto pyr by ∼9% (SST→Pyr connections from 12 to 10, and PV→Pyr connections from 17 to 15). In addition, the tonic inhibition of the Pyr neuron dendrites was reduced by 9%, and the conductance of excitatory OU processes on the Pyr neuron dendrites was reduced by 24%.

### False/failed signal detection rates

We calculated the error rates in stimulus detection by calculating the distribution of baseline firing rates of all Pyr neurons per circuit (sliding windows of 50 ms in 1 ms steps over 2 seconds pre-stimulus, for a total of 1950 windows), and the distribution of post-stimulus spike rates of all Pyr neurons in the 5 – 55 ms window post-stimulus across 100 randomized microcircuits (bootstrapped across all circuits, n = 1000 samples). The intersection point between the two distributions was set as the stimulus detection threshold. The probability of false detection was calculated by the integral of the pre-stimulus distribution above the detection threshold divided by the integral of the entire pre-stimulus distribution, and the probability of failed detection by the integral of the post-stimulus distribution below the detection threshold divided by the integral of the entire post-stimulus distribution.

## Statistical analysis

We calculated statistical significance using paired-sample or two-sample t-test (p < 0.05), where appropriate. We calculated the effect size using Cohen’s *d* (the difference in means divided by the pooled standard deviation).

## References

1. Lissemore, J. I. et al. Reduced GABAergic cortical inhibition in aging and depression. Neuropsychopharmacology 43, 2277–2284 (2018).

2. Northoff, G. & Sibille, E. Why are cortical GABA neurons relevant to internal focus in depression? A cross-level model linking cellular, biochemical and neural network findings. Mol. Psychiatry 19, 966–977 (2014).

3. Seney, M. L., Tripp, A., McCune, S., A. Lewis D., & Sibille, E. Laminar and cellular analyses of reduced somatostatin gene expression in the subgenual anterior cingulate cortex in major depression. Neurobiol. Dis. 73, 213–219 (2015).

4. Radley, J. J. et al. Repeated Stress Induces Dendritic Spine Loss in the Rat Medial Prefrontal Cortex. Cereb. Cortex 16, 313–320 (2006).

5. Radley, J. J. et al. Chronic behavioral stress induces apical dendritic reorganization in pyramidal neurons of the medial prefrontal cortex. Neuroscience 125, 1–6 (2004).

6. Bernardo, A. et al. Symptomatic and neurotrophic effects of GABAA receptor positive allosteric modulation in a mouse model of chronic stress. Neuropsychopharmacology 47, 1608–1619 (2022).

7. Kang, H. J. et al. Decreased expression of synapse-related genes and loss of synapses in major depressive disorder. Nat. Med. 18, 1413–1417 (2012).

8. Yao, H. K. et al. Reduced inhibition in depression impairs stimulus processing in human cortical microcircuits. Cell Rep. 38, (2022).

9. Guilloux, J.-P. et al. Molecular evidence for BDNF- and GABA-related dysfunctions in the amygdala of female subjects with major depression. Mol. Psychiatry 17, 1130–1142 (2012).

10. Lin, L. C. & Sibille, E. Somatostatin, neuronal vulnerability and behavioral emotionality. Mol. Psychiatry 20, 377–387 (2015).

11. Fee, C. et al. Behavioral deficits induced by somatostatin-positive GABA neuron silencing are rescued by alpha 5 GABA-A receptor potentiation. Int. J. Neuropsychopharmacol. (2021) doi:10.1093/ijnp/pyab002.

12. Prevot, T. D. et al. Reversal of Age-Related Neuronal Atrophy by α5-GABAA Receptor Positive Allosteric Modulation. Cereb. Cortex (2020) doi:10.1093/cercor/bhaa310.

13. Gidon, A. et al. Dendritic action potentials and computation in human layer 2/3 cortical neurons. Science 367, 83–87 (2020).

14. Beaulieu-Laroche, L. et al. Enhanced Dendritic Compartmentalization in Human Cortical Neurons. Cell 175, 643-651.e14 (2018).

15. Hay, E. & Segev, I. Dendritic Excitability and Gain Control in Recurrent Cortical Microcircuits. Cereb. Cortex N. Y. N 1991 25, 3561–3571 (2015).

16. Beniaguev, D., Segev, I. & London, M. Single cortical neurons as deep artificial neural networks. Neuron 109, 2727-2739.e3 (2021).

17. Liu, X. B., Zheng, Z. H., Xi, M. C. & Wu, C. P. Distribution of synapses on an intracellularly labeled small pyramidal neuron in the cat motor cortex. Anat. Embryol. (Berl.) 184, 313–318 (1991).

18. Chen, J. L. et al. Clustered dynamics of inhibitory synapses and dendritic spines in the adult neocortex. Neuron 74, 361–373 (2012).

19. Tsay, D. & Yuste, R. On the electrical function of dendritic spines. Trends Neurosci. 27, 77– 83 (2004).

20. Bekkers, J. & Hausser, M. Targeted dendrotomy reveals active and passive contributions of the dendritic tree to synaptic integration and neuronal output. Proc. Natl. Acad. Sci. U. S. A. 104, 11447–52 (2007).

21. Ichiyama, A. et al. State-dependent activity dynamics of hypothalamic stress effector neurons. eLife 11, e76832 (2022).

22. Larkum, M. E., Zhu, J. J. & Sakmann, B. Dendritic mechanisms underlying the coupling of the dendritic with the axonal action potential initiation zone of adult rat layer 5 pyramidal neurons. J. Physiol. 533, 447–466 (2001).

23. Rapp, M., Yarom, Y. & Segev, I. Modeling back propagating action potential in weakly excitable dendrites of neocortical pyramidal cells. Proc. Natl. Acad. Sci. 93, 11985–11990 (1996).

24. Gooch, H. M. et al. High-fidelity dendritic sodium spike generation in human layer 2/3 neocortical pyramidal neurons. Cell Rep. 41, 111500 (2022).

25. Lowe, G. Inhibition of backpropagating action potentials in mitral cell secondary dendrites. J. Neurophysiol. 88, 64–85 (2002).

26. Waters, J., Schaefer, A. & Sakmann, B. Backpropagating action potentials in neurones: measurement, mechanisms and potential functions. Prog. Biophys. Mol. Biol. 87, 145–170 (2005).

27. Testa-Silva, G. et al. High synaptic threshold for dendritic NMDA spike generation in human layer 2/3 pyramidal neurons. Cell Rep. 41, 111787 (2022).

28. Seeman, S. C. et al. Sparse recurrent excitatory connectivity in the microcircuit of the adult mouse and human cortex. eLife 7, e37349 (2018).

29. Obermayer, J. et al. Lateral inhibition by Martinotti interneurons is facilitated by cholinergic inputs in human and mouse neocortex. Nat. Commun. 9, (2018).

30. Molnár, G. et al. Human pyramidal to interneuron synapses are mediated by multi-vesicular release and multiple docked vesicles. eLife 5, e18167 (2016).

31. Hunt, S. et al. Strong and reliable synaptic communication between pyramidal neurons in adult human cerebral cortex. Cereb. Cortex bhac246 (2022) doi:10.1093/cercor/bhac246.

32. Destexhe, A., Rudolph, M., Fellous, J. M. & Sejnowski, T. J. Fluctuating synaptic conductances recreate in vivo-like activity in neocortical neurons. Neuroscience 107, 13–24 (2001).

33. Mazza, F., Guet-McCreight, A., Valiante, T. A., Griffiths, J. D. & Hay, E. In-silico EEG biomarkers of reduced inhibition in human cortical microcircuits in depression. PLOS Comput. Biol. 19, e1010986 (2023).

34. Guet-McCreight, A. et al. In-silico testing of new pharmacology for restoring inhibition and human cortical function in depression. Commun. Biol. 7, 1–13 (2024).

35. Salustri, C. et al. Cortical excitability and rest activity properties in patients with depression. J. Psychiatry Neurosci. 32, 259–266 (2007).

36. Okada, G., Okamoto, Y., Morinobu, S., Yamawaki, S. & Yokota, N. Attenuated Left Prefrontal Activation during a Verbal Fluency Task in Patients with Depression. Neuropsychobiology 47, 21–26 (2003).

37. Elliott, R. et al. Prefrontal dysfunction in depressed patients performing a complex planning task: a study using positron emission tomography. Psychol. Med. 27, 931–942 (1997).

38. Pu, S. et al. Reduced prefrontal cortex activation during the working memory task associated with poor social functioning in late-onset depression: Multi-channel near-infrared spectroscopy study. Psychiatry Res. Neuroimaging 203, 222–228 (2012).

39. Shestyuk, A. Y., Deldin, P. J., Brand, J. E. & Deveney, C. M. Reduced Sustained Brain Activity During Processing of Positive Emotional Stimuli in Major Depression. Biol. Psychiatry 57, 1089–1096 (2005).

40. Dorostkar, M. M., Zou, C., Blazquez-Llorca, L. & Herms, J. Analyzing dendritic spine pathology in Alzheimer’s disease: problems and opportunities. Acta Neuropathol. (Berl.) 130, 1–19 (2015).

41. Glantz, L. A. & Lewis, D. A. Decreased Dendritic Spine Density on Prefrontal Cortical Pyramidal Neurons in Schizophrenia. Arch. Gen. Psychiatry 57, 65–73 (2000).

42. Barch, D. M. et al. Selective Deficits in Prefrontal Cortex Function in Medication-Naive Patients With Schizophrenia. Arch. Gen. Psychiatry 58, 280–288 (2001).

43. Akram, A. et al. Stereologic estimates of total spinophilin-immunoreactive spine number in area 9 and the CA1 field: Relationship with the progression of Alzheimer’s disease. Neurobiol. Aging 29, 1296–1307 (2008).

44. Mineur, Y. S., Belzung, C. & Crusio, W. E. Effects of unpredictable chronic mild stress on anxiety and depression-like behavior in mice. Behav. Brain Res. 175, 43–50 (2006).

45. Dickstein, D. L., Weaver, C. M., Luebke, J. I. & Hof, P. R. Dendritic spine changes associated with normal aging. Neuroscience 251, 21–32 (2013).

46. Eyal, G. et al. Human Cortical Pyramidal Neurons: From Spines to Spikes via Models. Front. Cell. Neurosci. 12, (2018).

47. Yuste, R. Dendritic Spines and Distributed Circuits. Neuron 71, 772–781 (2011).

48. London, M. & Häusser, M. DENDRITIC COMPUTATION. Annu. Rev. Neurosci. 28, 503– 532 (2005).

49. Larkum, M. E. & Nevian, T. Synaptic clustering by dendritic signalling mechanisms. Curr. Opin. Neurobiol. 18, 321–331 (2008).

50. Golding, N. L., Kath, W. L. & Spruston, N. Dichotomy of Action-Potential Backpropagation in CA1 Pyramidal Neuron Dendrites. J. Neurophysiol. 86, 2998–3010 (2001).

51. Kuczewski, N. et al. Backpropagating Action Potentials Trigger Dendritic Release of BDNF during Spontaneous Network Activity. J. Neurosci. 28, 7013–7023 (2008).

52. Tripp, A. et al. Brain-Derived Neurotrophic Factor Signaling and Subgenual Anterior Cingulate Cortex Dysfunction in Major Depressive Disorder. Am. J. Psychiatry 169, 1194– 1202 (2012).

53. Oh, H. et al. The Role of Dendritic Brain-Derived Neurotrophic Factor Transcripts on Altered Inhibitory Circuitry in Depression. Biol. Psychiatry 85, 517–526 (2019).

54. Vigers, A. J. et al. Sustained expression of brain-derived neurotrophic factor is required for maintenance of dendritic spines and normal behavior. Neuroscience 212, 1–18 (2012).

55. Vetter, P., Roth, A. & Häusser, M. Propagation of Action Potentials in Dendrites Depends on Dendritic Morphology. J. Neurophysiol. 85, 926–937 (2001).

56. Hesprich, S. & Beardsley, S. Computational Characterization of the Cellular Origins of Electroencephalography. in 2019 9th International IEEE/EMBS Conference on Neural Engineering (NER) 352–355 (2019). doi:10.1109/NER.2019.8716883.

57. Reimann, M. W. et al. A Biophysically Detailed Model of Neocortical Local Field Potentials Predicts the Critical Role of Active Membrane Currents. Neuron 79, 375–390 (2013).

58. Van Geit, W. et al. BluePyOpt: Leveraging Open Source Software and Cloud Infrastructure to Optimise Model Parameters in Neuroscience. Front. Neuroinformatics 10, (2016).

59. Hay, E., Hill, S., Schürmann, F., Markram, H. & Segev, I. Models of Neocortical Layer 5b Pyramidal Cells Capturing a Wide Range of Dendritic and Perisomatic Active Properties. PLOS Comput. Biol. 7, e1002107 (2011).

60. Hay, E., Schürmann, F., Markram, H. & Segev, I. Preserving axosomatic spiking features despite diverse dendritic morphology. J. Neurophysiol. 109, 2972–2981 (2013).

61. 2010 Allen Institute for Brain Science. Allen Human Brain Atlas. Available from: human.brain-map.org.

62. Ponce, M. et al. Deploying a Top-100 Supercomputer for Large Parallel Workloads: the Niagara Supercomputer. in Proceedings of the Practice and Experience in Advanced Research Computing on Rise of the Machines (learning) 1–8 (Association for Computing Machinery, New York, NY, USA, 2019). doi:10.1145/3332186.3332195.

63. Mohan, H. et al. Dendritic and Axonal Architecture of Individual Pyramidal Neurons across Layers of Adult Human Neocortex. Cereb. Cortex 25, 4839–4853 (2015).

64. Fuhrmann, G., Segev, I., Markram, H. & Tsodyks, M. Coding of temporal information by activity-dependent synapses. J. Neurophysiol. 87, 140–148 (2002).

65. Mäki-Marttunen, T. et al. Alterations in Schizophrenia-Associated Genes Can Lead to Increased Power in Delta Oscillations. Cereb. Cortex N. Y. N 1991 29, 875–891 (2019).

66. Bryson, A. et al. GABA-mediated tonic inhibition differentially modulates gain in functional subtypes of cortical interneurons. Proc. Natl. Acad. Sci. 117, 3192–3202 (2020).

67. Moradi Chameh, H. et al. Diversity amongst human cortical pyramidal neurons revealed via their sag currents and frequency preferences. Nat. Commun. 12, 2497 (2021).

68. Scimemi, A. et al. Tonic GABAA receptor‐mediated currents in human brain. Eur. J. Neurosci. 24, 1157–1160 (2006).

69. Teleńczuk, B. et al. Local field potentials primarily reflect inhibitory neuron activity in human and monkey cortex. Sci. Rep. 7, 40211 (2017).

70. Gentet, L. J. et al. Unique functional properties of somatostatin-expressing GABAergic neurons in mouse barrel cortex. Nat. Neurosci. 15, 607–612 (2012).

71. Yu, J., Hu, H., Agmon, A. & Svoboda, K. Recruitment of GABAergic Interneurons in the Barrel Cortex during Active Tactile Behavior. Neuron 104, 412-427.e4 (2019).

72. Carnevale, N. T. & Hines, M. L. The NEURON Book. (Cambridge University Press, Cambridge, 2006). doi:10.1017/CBO9780511541612.

73. Hagen, E., Næss, S., Ness, T. V. & Einevoll, G. T. Multimodal Modeling of Neural Network Activity: Computing LFP, ECoG, EEG, and MEG Signals With LFPy 2.0. Front. Neuroinformatics 12, (2018).

74. Welch, P. The use of fast Fourier transform for the estimation of power spectra: A method based on time averaging over short, modified periodograms. IEEE Trans. Audio Electroacoustics 15, 70–73 (1967).

